# Drug screening linked to molecular profiling identifies novel dependencies in patient-derived primary cultures of paediatric high grade glioma and DIPG

**DOI:** 10.1101/2020.12.29.424674

**Authors:** Diana M Carvalho, Sara Temelso, Alan Mackay, Helen N Pemberton, Rebecca Rogers, Ketty Kessler, Elisa Izquierdo, Lynn Bjerke, Janat Fazal Salom, Matthew Clarke, Yura Grabovska, Anna Burford, Nagore Gene Olaciregui, Jessica KR Boult, Valeria Molinari, Mariama Fofana, Paula Proszek, Elisabet F Potente, Kathryn R Taylor, Christopher Chandler, Bassel Zebian, Ranj Bhangoo, Andrew J Martin, Bassam Dabbous, Simon Stapleton, Samantha Hettige, Lynley V Marshall, Fernando Carceller, Henry C Mandeville, Sucheta J Vaidya, Safa Al-Sarraj, Leslie R Bridges, Robert Johnston, Jane Cryan, Michael Farrell, Darach Crimmins, John Caird, Jane Pears, Giulia Pericoli, Evelina Miele, Angela Mastronuzzi, Franco Locatelli, Andrea Carai, Simon P Robinson, Mike Hubank, Michelle Monje, Andrew S Moore, Timothy EG Hassall, Angel Montero Carcaboso, Christopher J Lord, Mara Vinci, Chris Jones

## Abstract

Paediatric high grade glioma and diffuse midline glioma (including DIPG) are comprised of multiple biological and clinical subgroups, the majority of which urgently require novel therapies. Patient-derived *in vitro* primary cell cultures represent potentially useful tools for mechanistic and preclinical investigation based upon their retention of key features of tumour subgroups under experimental conditions amenable to high-throughput approaches. We present 17 novel primary cultures derived from patients in London, Dublin and Belfast, and together with cultures established or shared from Barcelona, Brisbane, Rome and Stanford, assembled a panel of 52 models under 2D (laminin matrix) and/or 3D (neurospheres) conditions, fully credentialed by phenotypic and molecular comparison to the original tumour sample (methylation BeadArray, panel/exome sequencing, RNAseq). In screening a subset of these against a panel of ~400 approved chemotherapeutics and small molecules, we identified specific dependencies associated with tumour subgroups and/or specific molecular markers. These included *MYCN*-amplified cells and ATM/DNA-PK inhibitors, and DIPGs with *PPM1D* activating truncating mutations and inhibitors of MDM2 or PARP1. Specific mutations in *PDGFRA* were found to confer sensitivity to a range of RTK inhibitors, though not all such mutations conferred sensitivity to targeted agents. Notably, dual PDGFRA/FGFR and downstream pathway MEK inhibitors showed profound effects against both PDGFRA-sensitising mutant and FGFR1-dependent non-brainstem pHGG and DIPG. In total, 85% cells were found to have at least one drug screening hit in short term assays linked to the underlying biology of the patient’s tumour, providing a rational approach for individualised clinical translation.

## INTRODUCTION

Paediatric high grade glioma (pHGG), including pontine and other diffuse midline glioma (DMG), is a remarkably heterogeneous collection of diseases, comprised of multiple biological subgroups arising in distinct anatomical locations within the central nervous system (CNS), and at different ages ^1^. In almost all instances, the clinical outcome remains extremely poor, with a median overall survival of 9-18 months ^2^. Making progress in this disease has been hampered historically by a lack of primary patient material for study, firstly in terms of defining the underlying biology, and latterly in terms of having appropriate model systems to develop novel therapeutic approaches. As progress has advanced rapidly in recent years to further refine our understanding of the critical genetic and epigenetic drivers of these diverse subgroups ^3–6^, it has become increasingly apparent that reappraisal of pHGG clinical trial structure is needed to evaluate novel agents in the appropriate patient populations ^7^, and that subgroup-specific models are required to generate the necessary preclinical evidence to prioritise novel therapies for trial ^8^. Given the rarity of the disease, this represents a significant challenge that can only be overcome collaboratively ^9^.

We published the first molecular characterisation of available cell line models of pHGG (n=3) more than a decade ago, before the advent of comprehensive genomic data on the tumours themselves ^10^. Grown as traditional monolayer cultures in serum-containing media, they represented amenable models given their rapid turnover *in vitro*, and ability to proliferate indefinitely, and with the later discovery of the histone H3 mutations which define a large proportion of pHGG/DIPG cases ^11,12^, even represented the first such model system available for a histone mutant tumour in the form of the H3.3G34V KNS42 cell line ^13^. Such models are however extremely limited, and concerns about how predictive they are of the patient disease have long been raised across all human cancer types ^14–16^ ^17,18^.

More recently, the establishment of short-term cultures in serum-free conditions directly from tumour cells collected at biopsy, surgery, or autopsy has allowed for the generation of a wealth of novel models across multiple cancer types. Such patient-derived (or ‘primary’) cell cultures are believed to be more reflective of the biology of the disease, given they preserve the genetic characteristics and heterogeneity of the tumours found in patients and could thus represent more clinically relevant models ^17,19–21^.

The first DIPG primary culture was established in 2011, in three-dimensional neurosphere conditions tailored to promote stem cell growth and restrict differentiation ^22,23^. Later, two-dimensional adherent culture of patient-derived glioma stem cell cultures was also employed 24. The experience in generating such cultures in DIPG specifically was consolidated in an international collaborative study identifying specific protocols predictive of success, including a source from biopsy *versus* autopsy, acquisition and transport in DMEM media *versus* Hibernate A, as laminin-coated 2D *versus* 3D neurospheres ^25^. Attempts to systematise the generation of and access to such models across childhood brain tumours in the form of collaborative biobanks have taken advantage of this increasing effort worldwide ^26^. Despite this, many of the pHGG/DIPG subgroups observed in the clinic remain poorly represented, and comprehensive molecular and phenotypic data on the models available is incomplete.

We have been prospectively collecting fresh tumour tissue to generate novel models from pHGG/DIPG patients treated at our hospitals, and have amassed a panel of 52 unique *in vitro* cultures, spanning multiple biological subgroups of the disease. Linking drug screening to comprehensive molecular analysis has identified numerous rational genetic dependencies that may be worthy of further investigation, and highlight the utility of such panels to inform future therapeutic development in this disease.

## RESULTS

### Establishment of novel pHGG/DIPG primary cultures

To facilitate the assembly of as large a panel of pHGG / DIPG cultures as possible, we took three approaches to sample collection (Figure 1A). We initiated prospective collection of fresh surgical material from the UK and Ireland; locally as part of the South Thames Multidisciplinary Paediatric Neuro-oncology Team (Kings College Hospital, St George’s Hospital, and the Royal Marsden Hospital NHS Trusts), and through the multidisciplinary services in Belfast (Royal Belfast Hospital for Sick Children) and Dublin (Beaumont Hospital, Temple Street Children’s University Hospital, and Our Lady’s Children’s Hospital). We were alerted to suspected high grade glial tumours from any anatomical site, with excess tumour material collected directly in Hibernate A transport medium for delivery to our labs in London, after a piece was frozen in liquid nitrogen, and along with blood for constitutional DNA. Up to June 2018, we prospectively collected 38 samples from 35 individual patients in this manner, 17 confirmed high grade gliomas, of which 14 were successfully established. A second source involved teams in Brisbane, Australia (through the Queensland Children’s Tumour Bank, n=5) and Rome, Italy (Ospedale Pediatrico Bambino Gesù, n=5) which had well-established collection procedures, and shipped us surgical material for culture as a ‘viable freeze’, *i.e.* tumour pieces minced and frozen slowly in DMSO. A total of 12 cultures received in this manner were subsequently established in our laboratory. Finally, we obtained 9 cultures from Barcelona, Spain (Hospital San Joan de Déu) and 15 from Stanford, USA (Stanford University) who had established cultures locally and shipped us vials of cells for expansion in our lab. Thus, in total we had a panel of 52 unique pHGG / DIPG cells for further analysis. All cell cultures were attempted in both two- laminin-coated flasks) and three- dimensions (neurospheres in ultra-low attachment plates) under stem cell conditions. Models were considered successfully established once they had reach passage five. Doubling times ranged from 1 to more than 6 days. Orthotopic implantation in the anatomical locations from which the primary tumour arose was attempted for 15 samples, of which 10 formed tumours with a median survival of 175 days, though with a wide variability amongst models (range 53-404 days) (Supplementary Figure S1). A summary of these data is provided in Supplementary Table S1.

**Figure 1.**
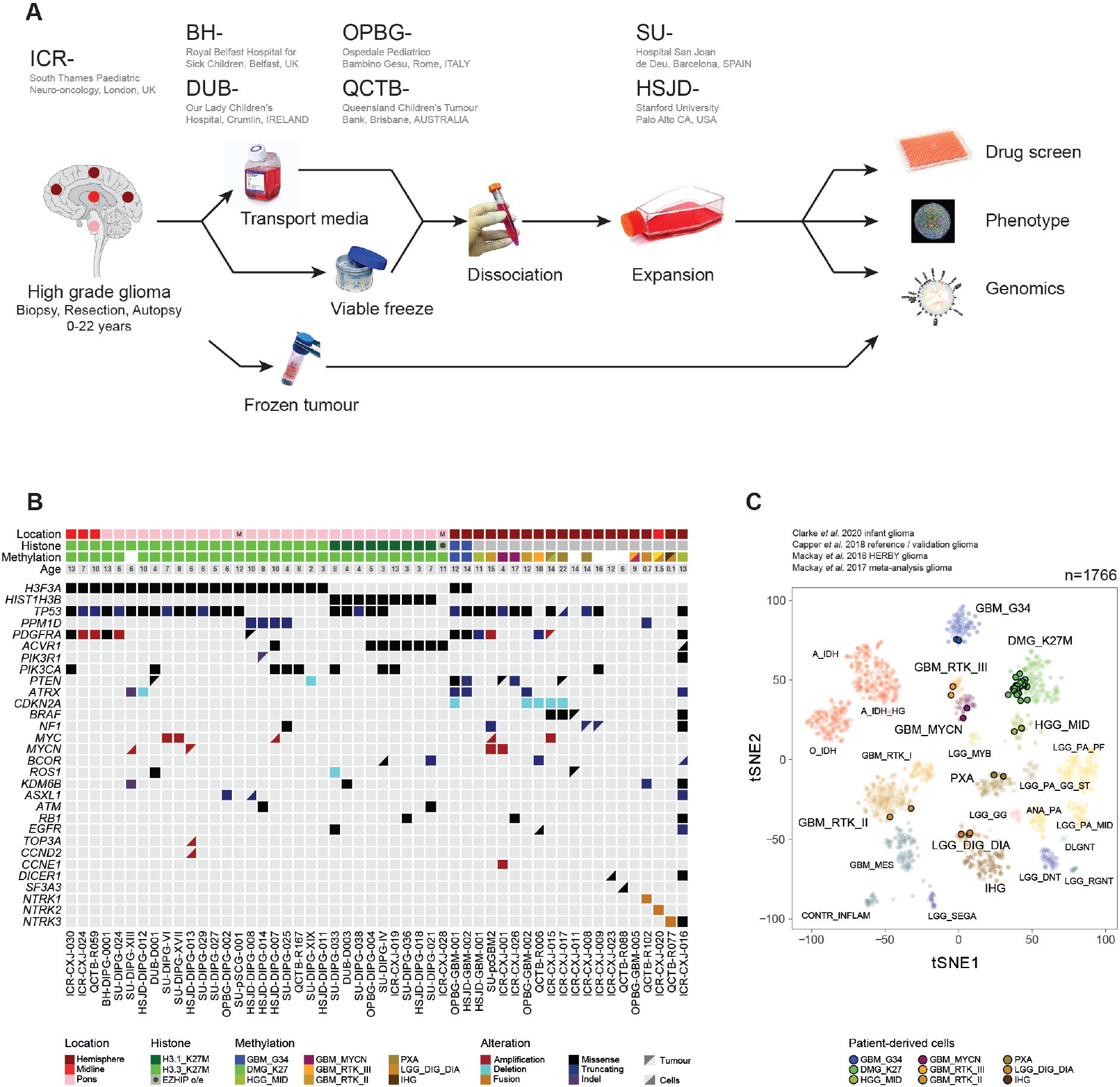
Patient-derived in vitro models of pHGG / DIPG. (A) Overview of study workflow. High grade glioma samples in patients under 23 years old are taken at biopsy, resection or autopsy. *In vitro* cell culture is established either fresh in transport medium, after shipping as viable cells, or from established lines. Cells are subjected to molecular analysis (along with frozen tumour, where possible), phenotypic characterisation and high-throughput drug screening. (B) Oncoprint representation of an integrated annotation of single nucleotide variants, DNA copy number changes and structural variants for all *in vitro* models (n=52). Samples are arranged in columns with genes labelled along rows. Clinicopathological and molecular annotations are provided as bars according to the included key. (C) t-statistic based stochastic neighbor embedding (t-SNE) projection of a combined methylation dataset comprising the *in vitro* models (circled) plus a reference set of glioma subtypes (n=1652). The first two projections are plotted on the × and y axes, with samples represented by dots colored by subtype according to the key provided.

### Molecular credentialing of established cells

Molecular analysis was carried out on cells once beyond at least passage five where possible. If available, analysis was also carried out on matched frozen tumour tissue (exome sequencing, RNAseq, methylation BeadArray) and blood (exome sequencing) from the same patient. Excluding one case identified as a hypermutator (ICR-CXJ-016, n=4043 somatic mutations in the tumour, n=4288 in the cells), we found an average of 24 and 19 alterations in the tumours and cells, respectively (Supplementary Table S2). There was no statistical difference in the number of alterations in tumours and cells (median = 24, range 6-229 and median = 19, range 5-163, respectively, p=0.645, t-test), with the vast majority of putative driver alterations, as defined by those most frequently observed in our large retrospective ^1^ and prospective ^8^ tumour series, found in both. These included 2 samples with *H3F3A* G34R, 22 with *H3F3A* K27M and 9 with *HIST1H3B* K27M mutations. There was one case of histone wild-type DIPG metastasizing to the spine, found instead to harbour *EZHIP* overexpression by RNAseq ^27^. We additionally had multiple samples with alterations in *TP53* (n=31), *PDGFRA* (n=11; 7 SNVs, 4 amplifications), *PIK3CA* (n=9), *ACVR1* (n=7), *ATRX* (n=5; 4 SNVs, 1 homozygous deletion), *PPM1D* (n=5), *CDKN2A/B* (n=5 homozygous deletions), *PTEN* (n=4; 3 SNVs, 1 homozygous deletion), *BRAF* (n=2), *NF1* (n=3), *BCOR* (n=3) and fusion genes involving *NTRK1/2/3* (n=3) (Figure 1B). Notable discrepancies included the acquisition of amplifications in *MYC/MYCN* and others in some cells (n=4) not observed in the tumour specimen sequenced, and a case of *BRAF*_600E present in the tumour, but not the cells (ICR-CXJ-011).

Using the Heidelberg methylation classifier ^28^ v11b4, cells (and where possible their matching tumour samples) were assigned scores according to which of 92 brain tumour subgroups they most closely resembled (Supplementary Table S2). In addition, the samples were clustered using a tSNE projection along with a reference pan-glioma set of 1766 tumour samples derived from the Heidelberg reference and validation sets ^28^ and our own studies ^1,8,29^. Although formal classifier scores for the cells was often equivocal, the primary cultures tended to align closely with both their relevant subgroups, as well as the patient-matched tumour specimens from which they were derived (Figure 1C). As such, our panel represented all major subgroups of pHGG/DIPG, with over half representing DMG_K27 (n=30), including the K27M wild-type DIPG spinal metastasis overexpressing *EZHIP*, and far fewer GBM_G34 (n=2). The remaining H3 wild-type hemispheric models represent HGG_MID (including the hypermutator case), GBM_MYCN, GBM_RTK_III, GBM_RTK_II as well as lower-grade subgroups such as PXA and the *NTRK*-fusion infant cases in the continuum between IHG and desmoplastic infantile ganglioglioma/astrocytoma (LGG_DIG_DIA) ^29^.

### High-throughput drug screening

For 20 of the cultures that we were able to establish as 3D neurospheres, we carried out 384 well plate drug sensitivity screening, using cell viability after five days drug exposure (estimated by CellTiter Glo) as the primary readout; this allowed us to calculate drug sensitivity Z scores, SF50 and AUC values for each drug in each 3D neurosphere culture (Figure 2A) (Supplementary Table S3). Where possible, two screens of FDA/EMA-approved drugs were performed, one where 80 drugs were screened at 8 different concentrations and a second screen profiling 397 drugs at 4 different concentrations (31 drugs were in both screens). Selected genotype-specific drug-sensitivity effects were validated in a secondary dose-response screen, where we used each drug at 12 different concentrations in appropriate sensitive and insensitive cell cultures.

**Figure 2.**
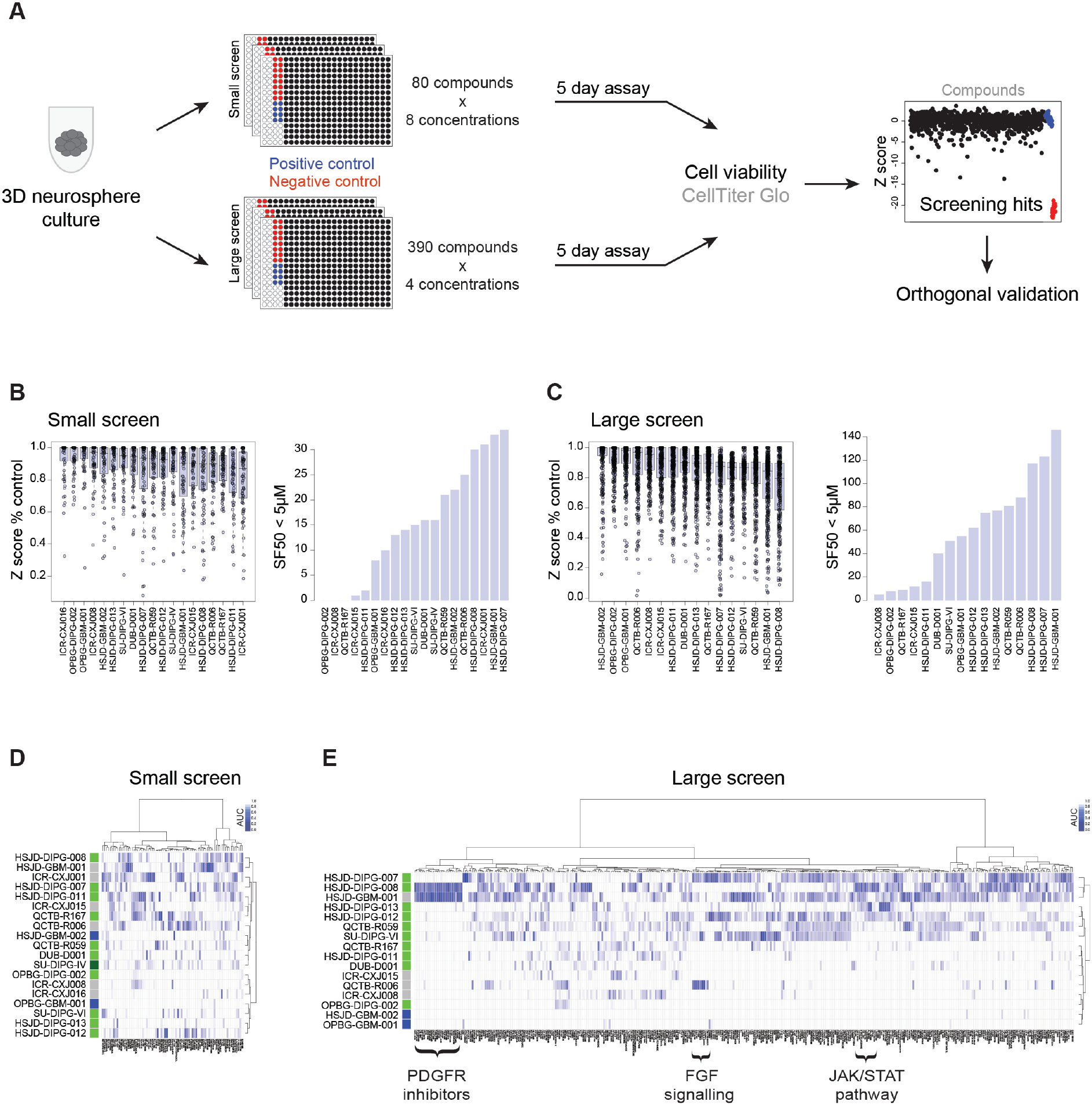
High-throughput drug screening of patient-derived cells. (A) Overview diagram of the drug screening process. Patient-derived cells (3D neurospheres) were seeded in 384-well plates in two screens of ~80 and ~400 compounds, with cell viability assessed after five days. Selected hits were validated by a full dose-response secondary screen. (B) Simple sensitivity metrics for the small screen. Left, boxplot of Z scores plotted as a percentage control (Z score POC). The thick line within the box is the median, the lower and upper limits of the boxes represent the first and third quartiles, and the whiskers 1.5x the interquartile range. Right, barplot of number of compounds for which the concentration of drug at which the surviving fraction of cells was <50% (SF50) was below 5μM. (C) Simple sensitivity metrics for the large screen. Left, boxplot of Z scores plotted as a percentage control (Z score POC). The thick line within the box is the median, the lower and upper limits of the boxes represent the first and third quartiles, and the whiskers 1.5x the interquartile range. Right, barplot of number of compounds for which the concentration of drug at which the surviving fraction of cells was <50% (SF50) was below 5μM. (D) Drug sensitivities in the small screen visualised by heatmap of area under the curve values (AUC), clustered by both drug (columns) and cells (rows). (E) Drug sensitivities in the large screen visualised by heatmap of area under the curve values (AUC), clustered by both drug (columns) and cells (rows).

Z scores calculated as a percentage of negative control (POC Z score) gave an indication of the relative effects on cell viability of each of the drugs in the screening panel, whilst the number of drugs for which the SF50 value was < 5μM gave a more direct assessment of the number of potential ‘hits’ for each cell model (Figure 2B,C). For the former, the median was 0.961-0.975, with a minimum of 0.128 (YM155), whilst for the latter this ranged from 5 drugs (ICR-CXJ-008) to 146 (HSJD-GBM-001), with a median of 76. There was no correlation of number of hits to *in vitro* doubling times in either the large (R^2^=0.111, p=0.245) or small screens (R^2^= 0.129, p=0.171, ANOVA).

Clustering the drug responses according to the AUC values across all cell lines highlighted common genetic dependencies shared by multiple cells with overlapping genotypes (Figure 2D,E). The most striking example was the large number of different multi-receptor tyrosine kinase inhibitors targeting PDGFRA in HSJD-GBM-001 and HSJD-DIPG-008, both of which were derived from tumours harbouring *PDGFRA* mutations. There were other conspicuous clusters of compounds targeting aspects of the JAK/STAT pathway in HSJD-DIPG-008 (JAK1/2, IKK, TORC1/2), and compounds targeting FGF signalling in QCTB-R006 (ERK, RAC, PPAR). Particularly for the large screen of nearly 400 compounds, there was clustering of the global response of the cells themselves on the basis of their genotype (H3 K27M, G34R and wild-type), with H3.3G34R cells specifically lacking in large numbers of drug sensitivity hits. Notably, the histone deacetylase (HDAC) inhibitor panobinostat, identified to be active in previous drug screens against DIPG cells ^30^, showed broad potency against a range of models (Supplementary Figure S2A). By contrast, the alkylating agent temozolomide, a commonly used chemotherapeutic in these diseases ^31^, showed no profound effects on cell viability (Supplementary Figure S2B).

### Single outlier responses

Certain cell models were found to be dramatically more sensitive to some classes of drugs compared to the rest of the panel. An exemplar of this was ICR-CXJ-001, which displayed screening hits on the basis of POC Z scores against a range of chemotypes of toolbox DNA repair inhibitors, specifically those targeting ATM (*e.g.* KU0060019) and DNA-PK (*e.g.* KU0057788) (Figure 3A). These cells were derived from a IDH wild-type grade IV glioblastoma, arising in the frontal lobe of a 4-year-old boy. The methylation subclassification was GBM_MYCN, and the DNA copy number profile from the methylation array revealed a high-level *MYCN* amplification (Figure 3B). This was confirmed by metaphase FISH analysis with specific *MYCN* and chromosome 2 centromeric probes, highlighting amplification via double minutes (Figure 3C). In validation, both compounds were found to have a significantly more potent effect on cell viability in ICR-CXJ-001 cells compared to a mini-panel of five pHGG/DIPG cells chosen as insensitive (and *MYCN* wild-type) (2.9-fold GI50, p<0.001, t-test) (Figure 3D), confirming the results of the initial screen. Similar outlier responses were observed in other cell models, such as DUB-D001 (H3.3K27M mutant DIPG), which was uniquely sensitive to inhibition of a senescence-associated axis of sphingosine 1-phosphate receptor (fingolimod) and Bcl-2/Bcl-xL (ABT-263, ABT-737) (Supplementary Table S3).

**Figure 3.**
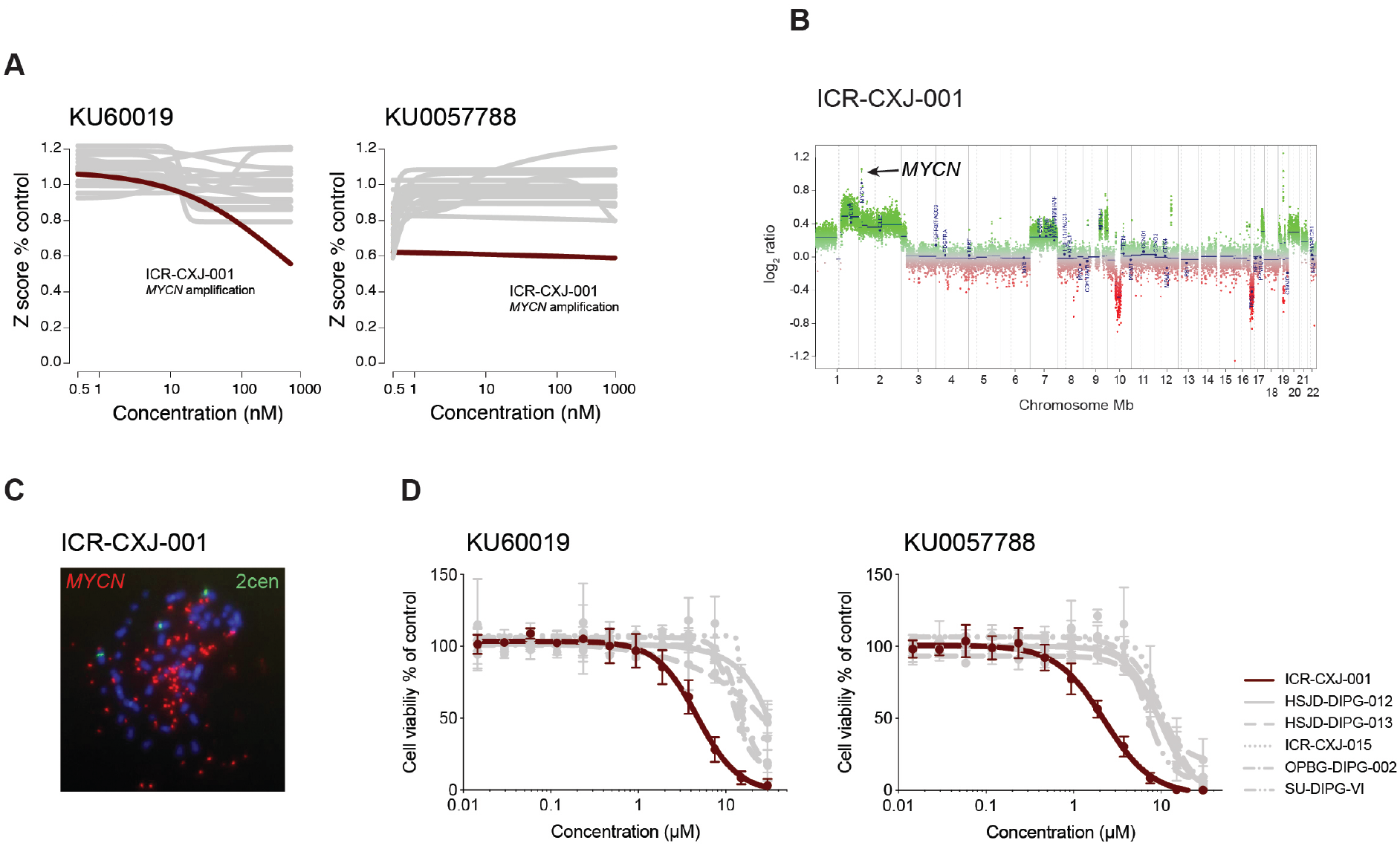
Single outlier screening hits. (A) Screen data for the ATM inhibitor KU60019 and the DNA-PK inhibitor KU0057788 in *MYCN*-amplified ICR-CXJ-001 cells (dark red) and the remaining panel of pHGG/DIPG cultures (grey). Concentration of compound is plotted on a log scale (x axis) against Z score plotted as a percentage of control (Z score POC) (y axis). (B) DNA copy number plot for ICR-CXJ-001 cells derived from methylation array data, with log2 ratios plotted (y axis) against genomic location by chromosome (x axis), and coloured green for gain, and red for loss. (C) Metaphase FISH carried out in ICR-CXJ-001 cells for *MYCN* (red) and chromosome 2 centromere (green). (D) Validation dose-response curves for the ATM inhibitor KU60019 and the DNA-PK inhibitor KU0057788 tested against *MYCN*-amplified ICR-CXJ-001 cells (dark red) and a mini-panel of pHGG/DIPG cultures (grey). Concentration of compound is plotted on a log scale (x axis) against cell viability (y axis). Mean plus standard error are plotted from at least n=3 experiments.

### Identification of genetic dependencies

We next sought to explore specific genetic dependencies across our panel by looking for correlations between drug class hits and pathway alterations in multiple cells. Strikingly, we observed consistent hits in drugs targeting the p53-mediated DNA damage response in cultures harbouring activating truncating mutations in *PPM1D* (Figure 4A). Three *PPM1D* mutant cultures (HSJD-DIPG-007, HSJD-DIPG-008, HSJD-DIPG-014) were significantly more sensitive on the basis of AUC to multiple, structurally distinct, PARP inhibitors (such as olaparib, p=0.0105, t-test), as well as the MDM2 inhibitor nutlin-3 (p=0.0165, t-test), compared to *PPM1D* wild-type cells (Figure 4B). This was also clearly seen with POC Z scores for both compounds (Figure 4C), and confirmed in dose-response validation experiments, with olaparib showing a 3.4-fold increased sensitivity (p=0.0103, t-test) (Supplementary Figure S3A), and nutlin-3 a 5.7-fold increased sensitivity (p<0.001, t-test) in *PPM1D* mutant compared to wild-type cells (Supplementary Figure S3B).

**Figure 4.**
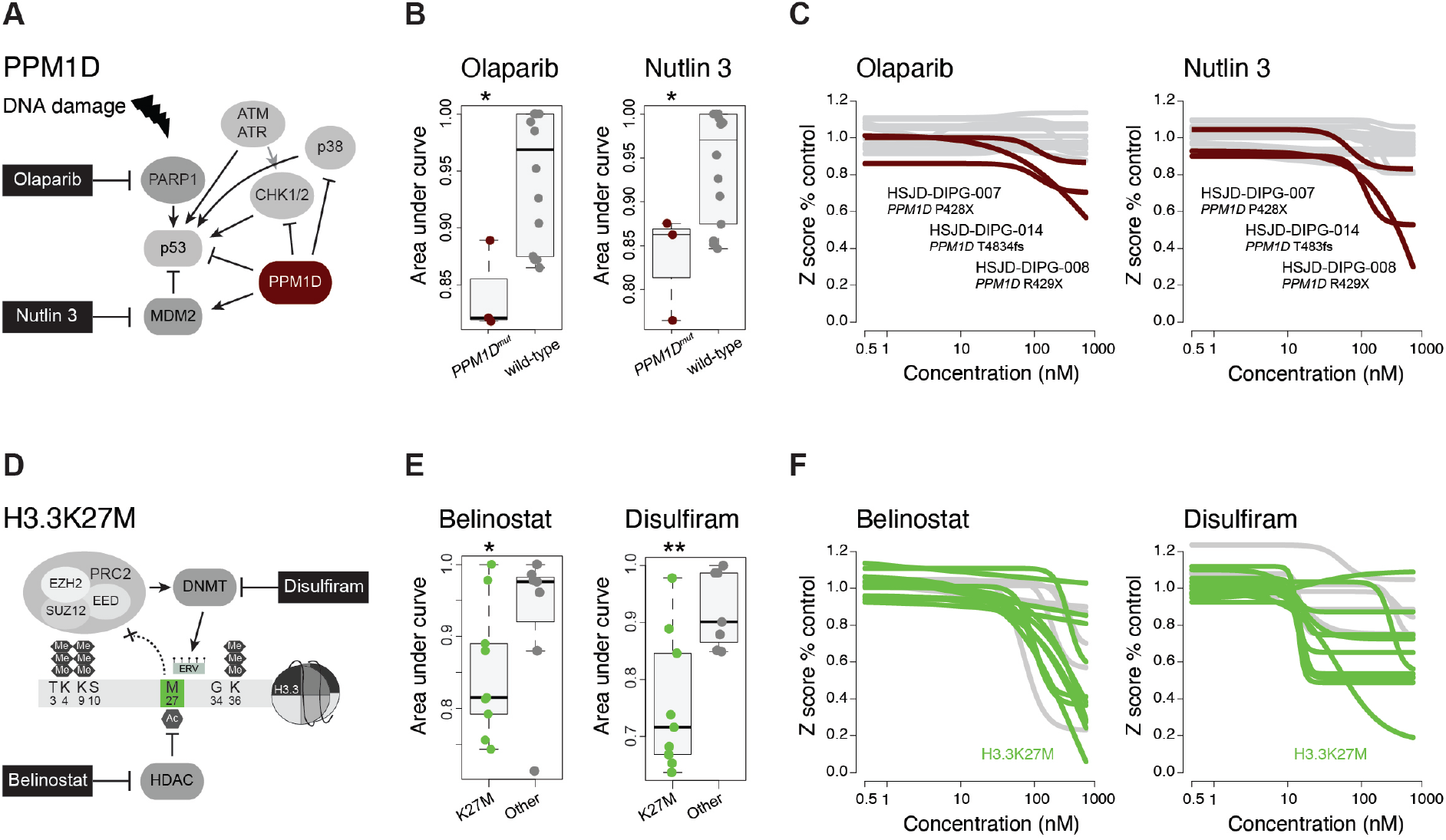
Genetic dependencies in pHGG/DIPG cells. (A) PPM1D pathway and control of the DNA damage response, showing critical nodes for which selective screening hits were seen (black boxes) in *PPM1D* mutant cells. (B) Boxplot of area under curve (AUC) values for the PARP inhibitor olaparib and the MDM2 inhibitor nutlin-3 separated by *PPM1D* status. The thick line within the box is the median, the lower and upper limits of the boxes represent the first and third quartiles, and the whiskers 1.5x the interquartile range. *p<0.05. (C) Screen data for olaparib and nutlin-3 in *PPM1D*-mutant cells (dark red) and the remaining panel of pHGG/DIPG cultures (grey). Concentration of compound is plotted on a log scale (x axis) against Z score plotted as a percentage of control (Z score POC) (y axis). (D) Histone H3.3 post-translational modifications and key interactors, showing critical nodes for which selective screening hits were seen (black boxes) in H3.3K27M mutant cells. (E) Boxplot of area under curve (AUC) values for the HDAC inhibitor belinostat and the DNMT inhibitor disulfiram, separated by H3.3K27M status. The thick line within the box is the median, the lower and upper limits of the boxes represent the first and third quartiles, and the whiskers 1.5x the interquartile range. **p<0.01, *p<0.05. (F) Screen data for belinostat and disulfiram in H3.3K27M-mutant cells (green) and the remaining panel of pHGG/DIPG cultures (grey). Concentration of compound is plotted on a log scale (x axis) against Z score plotted as a percentage of control (Z score POC) (y axis).

Similarly, H3.3K27M mutant cells were found to have differential sensitivities to compounds acting on the epigenetic processes dysregulated by the mutation. This included the well-established effects of HDAC inhibitors, in this instance most significantly by belinostat, but also a novel hit in the form of the acetaldehyde dehydrogenase inhibitor disulfiram, also thought to act on DNA methyltransferases (Figure 4D). There was a significantly increased sensitivity in H3.3K27M mutant cells compared to wild-type in terms of AUC (belinostat – p=0.0358, t-test; disulfiram – p=0.0088, t-test) (Figure 4E), as also seen with Z score POC (Figure 4F).

### Functional mutation annotation

We were also able to assign distinct and consistent responses to a range of similarly targeted agents to different genetic alterations targeting *PDGFRA* in our panel. Specifically, the two cultures showing a significantly different sensitivity to a wide range of compounds known to inhibit *PDGFRA* were derived from tumours harbouring A384ins (HSJD-GBM-001) and D846N (HSJD-DIPG-008) mutations, spanning both extracellular and kinase domains, respectively (Figure 5A). Conversely, cells harbouring the previously reported resistance mutation D842Y (HSJD-GBM-002) were insensitive as expected (Figure 5B) ^32^, however cells with additional *PDGFRA* mutations (including Y288C and K1061fs) or wild-type amplification were similarly lacking in response across a range of agents on the basis of Z score POC (Figure 5C). This was confirmed in dose-response validation, highlighting the exquisite sensitivity of the *PDGFRA*_A384ins HSJD-GBM-001 cells to crenolanib (Supplementary Figure S3C) and dasatinib (Supplementary Figure S3D) compared to other mutations, and the mini-panel of wild-type controls (6.0-fold, p=0.0035; 136-fold, p=0.0041, respectively, t-test). Notably, the D846N mutation in HSJD-DIPG-008 has been shown to be subclonal in multi-region autopsy sequencing of the patient’s tumour ^33^, and we were unable to detect this in the cell passages used for validation.

**Figure 5.**
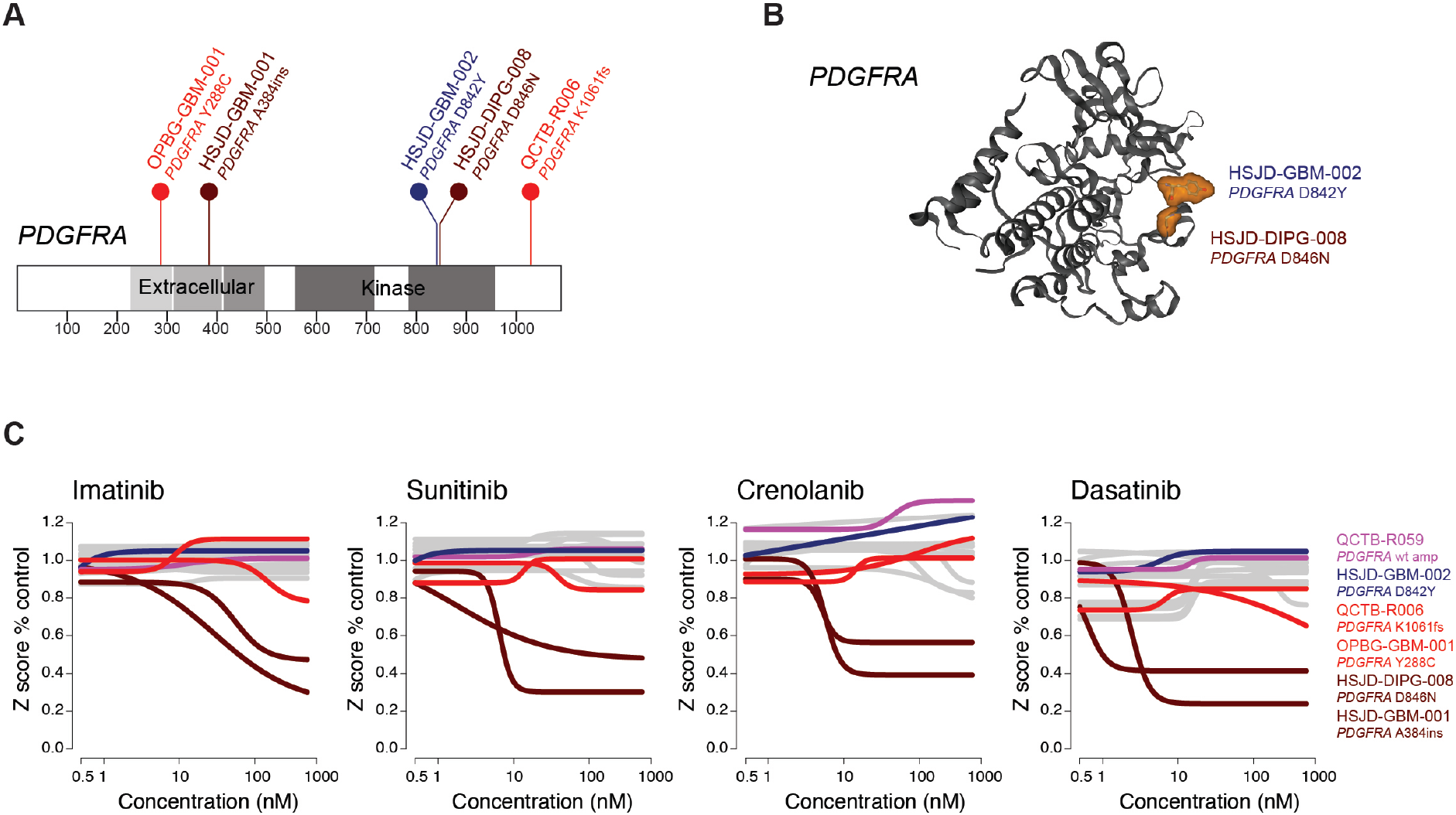
Functional annotation of PDGFRA mutations. (A) Cartoon representation of amino acid position of somatic *PDGFRA* mutations in five pHGG/DIPG samples, coloured by responsiveness to targeted inhibition (dark red, sensitive; red, insensitive; dark blue, resistant) with mutations found in *PDGFRA*, coloured by annotated functional domains and numbers provided for recurrent variants. Key functional domains are shaded. (B) Protein structure representation of PDGFRA showing mutant residues (shaded orange) for two closely located kinase domain mutations conferring sensitivity (D846N) and resistance (D842Y) to targeted inhibition, respectively. Generated in COSMIC-3D (cancer.sanger.ac.uk/cosmic3d). (C) Screen data for PDGFRA/multi-RTK inhibitors imatinib, sunitinib, crenolanib and dasatinib in cells associated with *PDGFRA* mutations (coloured according to responsiveness) and the remaining panel of pHGG/DIPG cultures (grey). *PDGFRA* wild-type amplified QCTB-R059 are in pink. Concentration of compound is plotted on a log scale (x axis) against Z score plotted as a percentage of control (Z score POC) (y axis).

### Pathway-level dependencies

Finally, we were additionally able to identify critical dependencies on multiple nodes of the same signalling pathway through our integrated drug screening and molecular data. Specifically, the RTK inhibitors ponatinib and PD173074 are known to inhibit both PDGFRA, and FGFR1, amongst other kinases ^34,35^ (Figure 6A). These drugs had significantly enhanced potency on the basis of Z score POC on HSJD-GBM-001 and HSJD-DIPG-008, but also on QCTB-R006 (Figure 6B), which harbours the C-terminal *PDGFRA* mutation K1061fs, and did not sensitise to the range of other PDGFRA inhibitors tested. This was confirmed by dose-response validation (ponatinib, 3.3-fold, p=0.0164, Supplementary Figure S3E; PD173074, 4.2-fold, p=0.0269, Supplementary Figure S3F, t-test), compared to the control mini-panel. These cells were clearly distinct from the others when their AUC values across all inhibitors reported to target FGFR1 were projected (Figure 6C).

**Figure 6.**
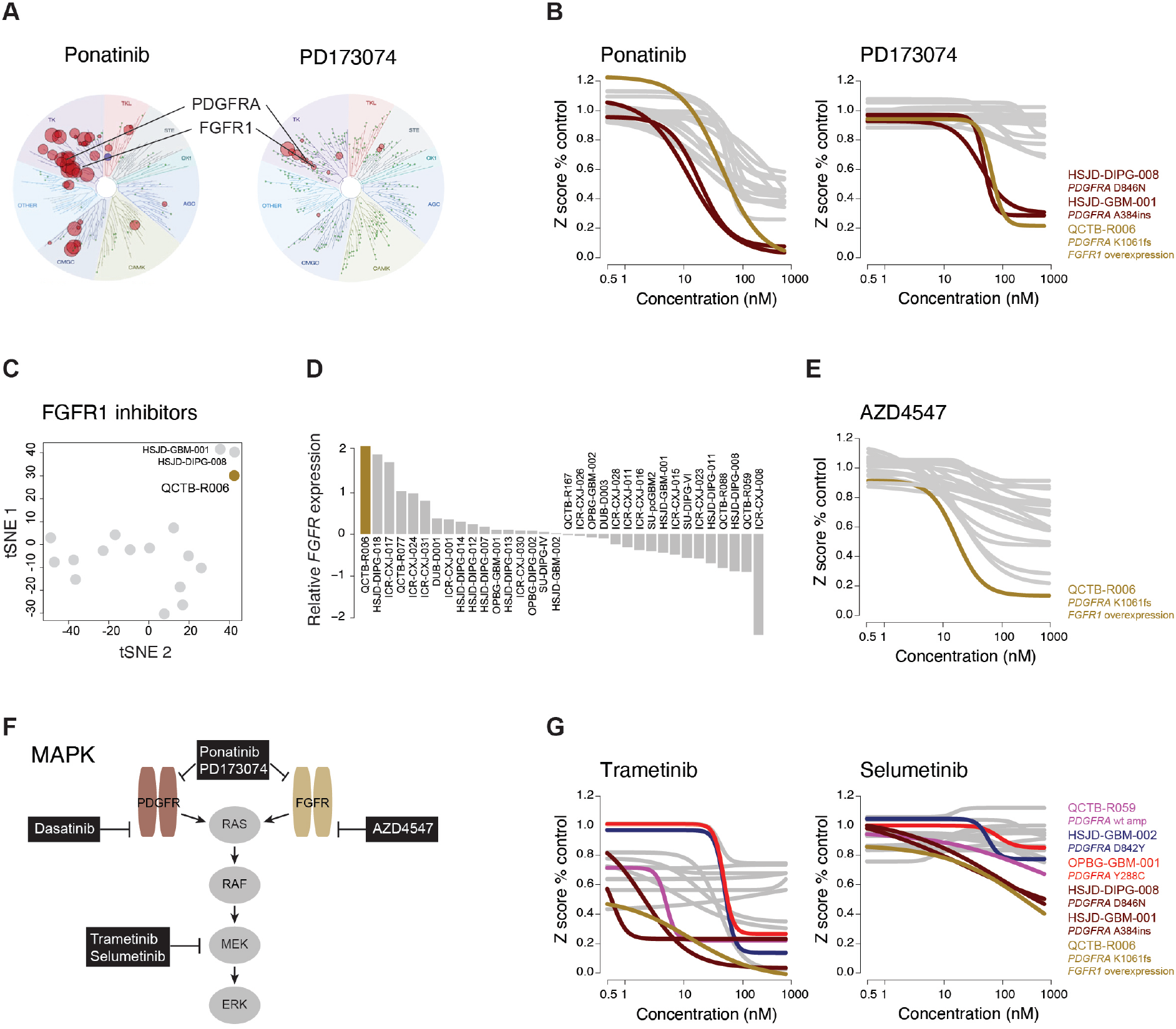
Pathway-level dependencies. (A) Kinome selectivity map for the multi-RTK inhibitors ponatinib and PD173074 at 10μM, with red circles reflecting kinases inhibited by >65%. PDGFRA and FGFR1 labelled. The diameter of the circles is inversely proportional to the percentage kinase activity remaining in the presence of inhibitor. Taken from the Harvard Medical LINCS KINOMEscan database (lincs.hms.harvard.edu). (B) Screen data for ponatinib and PD173074 in cells with sensitising *PDGFRA* mutation (dark red), *FGFR1*-overexpression (gold), and the remaining panel of pHGG/DIPG cultures (grey). Concentration of compound is plotted on a log scale (x axis) against Z score plotted as a percentage of control (Z score POC) (y axis). (C) tSNE projection of AUC values for all drugs in the screen known to target FGFR1, with QCTB-R006 highlighted in gold. (D) Barplot of *FGFR1* mRNA expression from RNA-seq counts of the panel of pHGG/DIPG cultures, ranked-ordered by relative expression. *FGFR1*-overexpressing QCTB-R006 cells are coloured gold. (E) Screen data for the FGFR1 inhibitor AZD4547 in cells with *FGFR1*-overexpression (gold), and the remaining panel of pHGG/DIPG cultures (grey). Concentration of compound is plotted on a log scale (x axis) against Z score plotted as a percentage of control (Z score POC) (y axis). (F) RTK/MAPK signalling pathway, showing critical nodes for which selective screening hits were seen (black boxes) associated with *PDGFRA* and/or *FGFR1* alterations/overexpression. (G) Screen data for the MEK inhibitors trametinib and selumetinib in cells with *FGFR1*-overexpression (gold), and distinct PDGFRA mutations (dark red, sensitive; red, insensitive; dark blue, resistant; pink, wild-type amplification), and the remaining panel of pHGG/DIPG cultures (grey). Concentration of compound is plotted on a log scale (x axis) against Z score plotted as a percentage of control (Z score POC) (y axis).

From RNA-seq data, QCTB-R006 cells had the highest levels of *FGFR1* mRNA expression in the entire panel, 4-fold above the median (Figure 6D). These cells were found to be differentially sensitive to the more specific FGFR1 inhibitor AZD4547, on the basis of Z score POC (Figure 6E), and confirmed by dose-response validation (7.5-fold, p=0.384, t-test) (Supplementary Figure S3G). With both PDGFRA and FGFR1 signalling through the MAPK pathway, we also explored the differential sensitivities of a range of downstream MEK inhibitors across our screening panel (Figure 6F). As examples, both trametinib and selumetinib were selectively potent against *PDGFRA*_A384ins HSJD-GBM-001, *PDGFRA*_D846N HSJD-DIPG-008, and *FGFR1*_overexpressing QCTB_R006 cells on the basis of Z score POC (Figure 6G). Notably, unlike that observed with RTK inhibitors, the *PDGFRA* wild-type amplified QCTB-R059 cells were also sensitive. These results were confirmed by dose-response validation for trametinib (112-fold, p<0.001, t-test), including the later passage HSJD-DIPG-008 in which we could not detect the *PDGFRA* mutation; importantly, also sensitive to MEK inhibition were the *BRAF*_V600E ICR-CXJ-015 cells (Supplementary Figure S3H). *PDGFRA*_D842Y HSJD-GBM-002 and *PDGFRA*_Y288C OPBG-GBM-001 cells remained insensitive.

## DISCUSSION

The establishment of methods for generating patient-derived cultures of pHGG and DIPG is providing a rapidly expanding resource for the study of the disease, with such models being used for mechanistic evaluation of epigenetic reprograming associated with the histone H3 mutations ^36–40^, tumour-tumour cell ^33^ and tumour-microenvironmental interactions ^41,42^, invasion/migration ^43^ and the evaluation of novel drug targets by preclinical efficacy assays ^44–50^ and high-throughput screening ^30,51,52^. Here we present our experience with deriving a new, well-characterised, prospectively-collected panel of such models alongside more established cells, in order to inform such initiatives spanning pHGG/DIPG subgroups. In addition, we demonstrate the utility of a subset of these models to identify novel, or refined, candidates for biologically rational therapeutic approaches.

We present around 40 models that have not previously had their fundamental molecular features published, though many of which have already been shared collaboratively across research groups worldwide. These cells present a wide range of behaviours *in vitro*, in both 2D and 3D serum-free culture, and many are problematic to include in high-throughput approaches. Given the additional practical difficulties associated with protracted latency times when orthotopically implanted in immunocompromised mice, often between 12-18 months in our hands, we do not yet have systematic *in vivo* characterisation of the full panel, particularly given the criteria of deriving a serially xenografted P2 model in order to consider a PDX model ‘established’ ^26^. Such efforts are, however, ongoing. These data are important to record and make available, so that researchers are clear about which models may be most suitable for their particular experiments, in addition to the key molecular features they wish to explore. Where models have been successful, *in vitro* and *in vivo*, they can be seen to recapitulate key phenotypic features of the human disease in terms of morphology, marker expression ^33^, and infiltrative growth.

An important criterion to assess the usefulness of these models is how well they retain the key molecular features of the human disease in general, and specifically the patient-matched tumour samples from which they were derived. Here we show that the major driving alterations are retained, and that epigenetic profiles of cell-based models closely resemble the tumours themselves. These data can be reliably integrated to match these models to emerging and well-established subtypes of pHGG/DIPG ^1^. These include models of a number of subtypes previously poorly represented, including H3.1K27M DIPG (n=9), non-brainstem H3.3K27M DMG (n=3), and a K27M wild-type DIPG (spinal metastasis) with EZHIP overexpression ^27^, in addition to multiple hemispheric subtypes and infantile hemispheric glioma with *NTRK* fusions (n=3). Many remain in only limited numbers, in particular H3.3G34R/V, GBM_MYCN etc, though we are aware they are beginning to emerge in several centres ^26^.

By applying a subset of the most amenable models to high-throughput drug screening, we have been able to identify novel treatment candidates via a variety of means. This includes the systematic assessment of genotype-phenotype correlations in the form of drug sensitivities linked to specific genetic alterations, and can be done at the gene- or pathway-level. Hits identified this way included the plausible synthetic lethalities of drugs targeting p53-mediated DNA repair in DIPG patients with activating truncating mutations in *PPM1D*. Such tumours represent ~10% DIPG ^3–6^ and 37.5% adult midline glioma ^53^, and have been associated with hypersensitivity to NAMPT inhibitors ^54^, or a direct target for drug inhibition in combination with radiotherapy ^55^. PPM1D (or Wip1) is described as a negative regulator of the p53-mediated DNA damage response ^56^, and somatic mutations are mutually exclusive with those found in *TP53* ^1^. Identification of hypersensitivity to MDM2 and PARP inhibitors for this genotype opens up the possibility of refinement of patient stratification for future clinical trials of these agents in pHGG/DIPG.

An unexpected finding from this approach was the differential sensitivity of disulfiram, marketed as Antabuse in the treatment of chronic alcoholism ^57^, in H3.3K27M cells. Although, primarily an inhibitor of acetaldehyde dehydrogenase, reports have suggested disulfiram to also have numerous additional effects on diverse processes such as the proteasome, PLK1 and NF-KB signalling ^58^. Intriguingly, its copper-containing metabolite CuET (which spontaneously forms in cell culture media) is thought to kill cancer cells through aggregation of the p97/VCP segregase subunit NPL4 ^59,60^, which removes ubiquitinated proteins from the chromatin fraction and may indicate a more direct effect on histones in K27M cells. It has also been shown to be a non-nucleoside DNMT1 inhibitor that can reduce global ^5me^C content, and reactivate epigenetically silenced genes in prostate cancer cells ^61^. Such agents, like 5-azacytidine, have been suggested as treatments for H3K27M cells owing to the observed pervasive H3K27ac deposition across multiple loci including endogenous retroviral elements (ERVs), priming them for activation by DNA methylation, and rendering them differentially sensitive to DNA demethylating agents ^39^. Notably, the combination of such drugs with HDAC inhibitors presented a specific therapeutic vulnerability in H3K27M cells, and repurposed agents such as belinostat and disulfiram may contribute to such approaches.

By contrast, it is apparent that not all genetic alterations in any given gene necessarily convey the same level of sensitivity to targeted inhibition. Numerous PDGFRA inhibitors have been trialled in pHGG/DIPG since the earliest molecular profiling studies identifying a relatively high frequency of amplification and mutation ^62,63^. The results have been generally / universally disappointing, in part through likely poor blood-brain barrier penetration of certain compounds, and also due to a lack of appropriate patient selection ^64^. We show this is compounded by the fact that certain mutations (and amplification of the wild-type) may be non-responsive to RTK inhibition.

It does also represent an opportunity, whereby patient-derived models can be used to determine the relative sensitivity to these agents of the distinct mutations – either through retrospective screening panels like this one, or through co-clinical trials ^65^. Given the rarity of the disease, such variant-level sensitivities may pose a problem for running clinical trials, however our data also highlight the utility of identifying specific downstream pathway vulnerabilities, and/or multi-target inhibitors in which common susceptibilities to drugs may be identified across genotypes – the example here whereby patients with both brainstem and non-brainstem tumours with sensitizing alterations in *PDGFRA* or *FGFR1*, who may benefit from dual RTK or MEK inhibitors and potentially be included in the same study.

There are now many initiatives worldwide to develop, characterise and utilise such patient-derived models of pHGG / DIPG. Aggregating ongoing efforts, even if only virtually, to provide the most appropriate, well-characterised models in a systematic way to adequately resource the whole research community remains a priority.

## MATERIALS and METHODS

### Patient samples

From the South Thames paediatric neurosurgical centres (Kings College Hospital and St George’s Hospital NHS Trusts), where the oncology care is delivered at the Royal Marsden Hospital Children and Young People’s Unit, we collected tumour tissue directly from theatre for any suspected high grade glial tumours from patients under the age of 25 years at first diagnosis. These were taken from any anatomical site, and whenever possible, as excess to routine diagnostics, were collected from biopsies as well as resections. If the pathological diagnosis was not a WHO grade III or IV glioma, the specimen was banked for any future appropriate Ethical Committee-approved project. The median age of the high grade glioma cases from which we successfully established cell cultures was 13.0 years (range 1.5 – 22.0), and included 11 boys and 3 girls. These samples and additional DIPG specimens from Dublin were collected in Hibernate A transport medium (Thermo Fisher Scientific, Waltham MA, USA, A1247501) and processed further in our laboratory. Further prospectively collected samples were instead shipped as live minced cryopreserved tissue in Dulbecco’s Modified Eagles Medium: Nutrient Mixture F12 (DMEM/F12; Thermo Fisher Scientific, 11330-038) supplemented with 10% DMSO and 0.1% Bovine Serum Albumin (BSA). Where possible, samples of fresh-frozen tissue and blood were also provided for each case. All patient samples were collected under full Research Ethics Committee approval at each participating centre. A summary of these data is provided in Supplementary Table S1.

### Nucleic acid extraction

DNA and RNA were extracted following the DNeasy Blood & Tissue kit (QIAGEN, Hilden, Germany, 69504) and the RNeasy Plus Mini Kit protocols (QIAGEN, 74134), respectively. Occasionally, a dual RNA/DNA extraction kit, Quick-DNA/RNA™ Miniprep Plus Kit (Zymo Research, Irvine CA, USA, D7003T), was also used following manufacturer’s instructions. Concentrations were measured using the Qubit dsDNA Assay Kits (Thermo Fisher Scientific, Q32850, Q32851) and/or a TapeStation 4200 (Agilent, Santa Clara CA, USA).

### Primary cell cultures

DIPG and pGBM patient-derived cultures established in our lab and others were grown in stem cell media consisting of 250ml DMEM/F12 (Thermo Fisher Scientific, 11330-038), 250ml Neurobasal-A Medium (Thermo Fisher Scientific, 10888-022), 10mM HEPES Buffer Solution (Thermo Fisher Scientific, 15630-080), 1mM MEM Sodium Pyruvate Solution (Thermo Fisher Scientific, 11360-070), 0.1 mM MEM Non-Essential Amino Acids Solution (Thermo Fisher Scientific, 11140-050) and 1x Glutamax-I Supplement (Thermo Fisher Scientific, 35050-061). The media was supplemented with B-27 Supplement Minus Vitamin A 1:50 (Thermo Fisher Scientific, 12587-010), 20ng/ml recombinant Human-EGF (2B Scientific LTD, Oxford, UK, 100-26), 20ng/ml recombinant Human-FGF (2B Scientific LTD, 100-146), 10ng/ml recombinant Human-PDGF-AA (2B Scientific LTD, 100-16), 10ng/ml recombinant Human-PDGF-BB (2B Scientific LTD, 100-18), and 2μg/ml Heparin Solution (Stem Cell Technologies, Cambridge, UK, 07980).

Patient-derived cultures were established either immediately after collection (biopsy, resection or autopsy) or from live cryopreserved tissue, with authenticity verified using short tandem repeat (STR) DNA fingerprinting ^22,30^ (Supplementary Table S2) and certified mycoplasma-free. Fresh tumour tissue was first finely minced with the use of sterile scalpels followed by gentle enzymatic dissociation with LiberaseTL (Roche, Basel, Switzerland, 5401020001) for 10 min at 37°C. After incubation, an additional 5ml of fresh media were added and the sample was centrifuged at 1300rpm for 5 minutes. The digested tissue was then resuspended in fresh media and triturated gently with a pipette (5-10 times). The cell suspension was then transferred into a laminin coated flask (2D, two-dimensional culture) (Millipore, Burlington MA, USA, CC095) and/or a ultra-low attachment flask (Corning, 3814) (3D, three-dimensional culture). Infant samples were transferred into laminin/fibronectin coated flasks. When necessary, red blood cells were lysed by using the ACK lysis buffer (Thermo Fisher Scientific, A1049201). Cells were incubated at 37°C, 5% CO2, 95% humidity. Cells were passaged into new flasks when cultures reached confluence (90% surface area coverage for laminin cells and a diameter of 200μm for 3D cultures). Neurospheres were centrifuged in their original media at 900rpm for 10 minutes, and the pellet resuspended and incubated for 3-7 minutes at 37°C in accutase dissociation reagent (Sigma, Poole, UK, A6964). After enzyme neutralization with fresh medium, cells were centrifuged at 1300rpm for 5 minutes. The pellet was resuspended in 200ul fresh media to generate a single cell suspension by pipetting up and down for 30-50X. To passage the laminin adherent cells, the medium was removed from the flask and cells were incubated with accutase for 2-5 minutes at 37°C. Once the cells were detached, media was added and transferred to a universal tube for centrifugation at 1300rpm for 5 minutes. The pellet was then resuspended in fresh media.

Doubling times for 2D and 3D patient-derived lines were estimated by seeding 1000-5000 cells/well in black 96 well plates and cell viability was measured with CellTiter-Glo (Promega, Southampton, UK, G7571/G9682) every 2-3 days. Values were plotted and the exponential part of the curves was used to calculate the doubling times following the equation

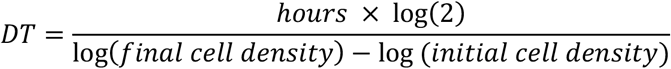

### FISH

Briefly, colcemid was added to media containing cell suspensions and left overnight. They were then incubated with 0.075M KCL at 37ºC for 5 – 30 minutes before being fixed with ice-cold methanol and acetic acid (3:1 ratio) and dropped from height onto a slide. The MYCN probe (VYSIS LSI N-MYC 2P24.1 Spectrum Orange) was purchased from Abbott Molecular (Des Plaines, IL, USA). BAC clones mapping to CEP2 (RP11-685C07) were purchased from BACPAC Resources Center (Children’s Hospital Oakland Research Institute, Oakland, CA, USA). BAC DNA was amplified using the IllustraTM GenomiPhiTM V2 DNA amplification kit (GE Healthcare, Little Chalfont, UK) and probes were labelled with DIG using Digoxigenin-11-dUTP (Roche, Burgess Hill, UK) and the BioPrime® DNA Labeling System (Invitrogen, Paisley, UK) according to the manufacturer’s instructions. Slides were treated with 70% acetic acid digested in 0.025% pepsin (Sigma) in 0.01M HCL at 37°C for 5 minutes before adding the prepared probes with (75%) hybridisation mix under a coverslip and denatured for 5 minutes at 73°C on a heat block. Slides were then incubated overnight in a humidified chamber at 37°C, mounted in Vectashield with DAPI (Vector Laboratories, Peterborough, UK), and captured on the Metasystems (Altlussheim, Germany) Axioskop fluorescence microscope (Zeiss, Cambridge, UK) using filters for DAPI, Cy3 and FITC.

### Methylation profiling

Methylation analysis was performed using either Illumina 450K or EPIC BeadArrays at University College London (UCL) Great Ormond Street Institute of Child Health, Bart’s Cancer Centre, Bambino Gesù Children’s Hospital, or DKFZ Heidelberg. Data was pre-processed using the minfi package in R (v11b4). DNA copy number was recovered from combined intensities using the conumee package. The Heidelberg brain tumour classifier (molecularneuropathology.org) ^28^ was used to assign a calibrated score to each case, associating it with one of the 91 tumour entities which feature within the current classifier. Clustering of beta values from methylation arrays was performed based upon correlation distance using a ward algorithm. DNA copy number was derived from combined log2 intensity data based upon an internal median processed using the R packages minfi and conumee to call copy number in 15,431 bins across the genome.

### DNA and RNA sequencing

DNA was sequenced either as whole genome or captured using Agilent SureSelect whole exome v6, xGen Exome Research panel v1 (Integrated DNA Technologies, Leuven, Belgium), or a custom panel of 330 genes known to present in an unselected series of pHGG 1. Libraries were prepared from 50-200 ng of DNA using the Kapa HyperPlus kit and DNA was indexed utilising 8bp-TruSeq-Custom Unique Dual Index Adapters (Integrated DNA Technologies). Following fragmentation, DNA was end-repaired, A-tailed and indexed adapters ligated. DNA was amplified, multiplexed and hybridized using 1 μg of total pre-capture library. After hybridization, capture libraries were amplified and sequencing was performed on either a MiSeq, NextSeq500 or NovaSeq6000 system (Illumina) with 2 × 150bp, paired-end reads following manufacturer’s instructions.

Ribosomal RNA was depleted from 150-2000 ng of total RNA from fresh frozen (FF) and formalin fixed paraffin-embedded (FFPE) tissue using NEBNext rRNA Depletion Kit. Following First strand synthesis and directional second strand synthesis resulting cDNAs were used for library preparation using NEBNext Ultra II Directional RNA library prep kit for Illumina performed as per the manufacturer’s recommendations. Exome capture reads were aligned to the hg19 build of the human genome using bwa v0.7.12 (bio-bwa.sourceforge.net), and PCR duplicates removed with PicardTools 1.94 (pcard.sourceforge.net). Single nucleotide variants were called using the Genome Analysis Tool Kit v3.4-46 based upon current Best Practices using local re-alignment around indels, downsampling and base recalibration with variants called by the Unified Genotyper (broadinstitute.org/gatk/). Variants were annotated using the Ensembl Variant Effect Predictor v74 (ensembl.org/info/docs/variation/vep) incorporating SIFT (sift.jcvi.org) and PolyPhen (genetics.bwh.harvard.edu/pph2) predictions, COSMIC v64 (sanger.ac.uk/ genetics/CGP/cosmic/), dbSNP build 137 (ncbi.nlm.nih.gov/sites/SNP), ExAc and ANNOVAR annotations. RNA sequences were aligned to hg19 and organized into de-novo spliced alignments using bowtie2 and TopHat version 2.1.0 (ccb.jhu.edu/software/tophat). Fusion transcripts were detected using chimerascan version 0.4.5a filtered to remove common false positives.

### Drug screen

A minimum of 1.75 × 10^7^ cells were screened in 3D in 384-well flat-bottom plates containing eight concentrations (0.5, 1, 5, 10, 50, 100, 500, 1000 nM) of each of the 397 FDA/EMA-approved drugs assayed. Cells were seeded at 1500 cells per well and continuously cultured in the presence of drugs for five days, after which cell viability was estimated by the use of 3D CellTitre-Glo reagent (Promega, G9682). Luminescence values from each well in the 384 well plate were normalised to the median of signals from wells containing cells exposed to the drug vehicle, DMSO, generating Surviving Fractions (SF). High-throughput drug screening data was also pre-processed and normalised with the cellHTS2 package in R to produce dose-response curves and Z-scores. R-package drc was used to calculate area under the curve (AUC) values from dose-response curves. AUC values for each drug in each cell line were standardised generating Z scores to allow for comparisons to be made between cell lines.

### Validation

Small molecules were purchased as solid from Selleckchem (Houston TX, USA) with the exception of TTP22 which was purchased from APExBIO (Houston TX, USA) and stored in DMSO at 10mM stocks. Prior to screening, the compounds and DMSO controls were arrayed in V-Bottom 384-well plates (Greiner, Kremsmuenster, Austria, 781280) using the ECHO 550 liquid dispenser (Labcyte, San Jose CA, USA) at 12 different concentrations. Cells were plated in 384-well plates (Greiner, 781090) at 1500 cells per well in 40μl media and allowed to grow for 24h. On day 2, 50μl of media was added per well in order to resuspend the compounds to 5X concentration. 10μl of media containing the compounds was then transferred to the 384-well plate containing the cells. After 5 days of incubation, cell viability was measured using 3D CellTiter–Glo (Promega, G9682). A minimum of 3 technical replicates was used per drug concentration and at least 3 biological replicates were performed. GI50 values were calculated using GraphPad Prism version 8 as the concentration of compound required to reduce cell viability by 50%.

### In vivo

For intracranial implantation, all experiments were performed in accordance with the local ethical review panel, the UK Home Office Animals (Scientific Procedures) Act 1986, the United Kingdom National Cancer Research Institute guidelines for the welfare of animals in cancer research and the ARRIVE (Animal Research: Reporting *In Vivo* Experiments) guidelines ^66,67^. Single cell suspensions were obtained immediately prior to implantation in either NOD.Cg-*Prkdc^scid^ Il2rg^tm1WjI^*/SzJ (NSG), NOD.Cg-*Prkdc^scid^*/J (NOD.SCID) or *Foxn1^nu^* (Athymic Nude) mice (Charles River, Harlow, UK). Animals were anaesthetized with 4% isoflurane and maintained at 2-3% isoflurane delivered in oxygen (1L/min). Core body temperature was maintained using a thermo-regulated heated blanket. A subcutaneous injection of buprenorphine (0.03mg/Kg) and Meloxicam (5mg/Kg) was given for general analgesia. Animals were depilated at the incision site and Emla cream 5% (lidocaine/prilocaine) was applied on the skin. The cranium was exposed via midline incision under aseptic conditions, and a 31-gauge burr hole drilled above the injection site. Mice were then placed on a stereotactic apparatus for orthotopic implantation. The coordinates used for the cortex were x=−2.0, z=+1.0, y=−2.5mm from the bregma, for the thalamus x=−1.2, z=2, y=3mm from the bregma, and for the pons x=+1.0, z=−0.8, y=−4mm from the lambda. 250,000-500,000 cells in 2-7μL were stereotactically implanted using a 25-gauge SGE standard fixed needle syringe (SGE™ 005000) at a rate of 2μl/min using a digital pump (HA1100, Pico Plus Elite, Harvard Apparatus, Holliston, MA, USA). At the completion of infusion, the syringe needle was allowed to remain in place for at least 2 minutes, and then manually withdrawn slowly to minimize backflow of the injected cell suspension. Mice were monitored until fully recovered from surgery. 24h post-surgery a subcutaneous injection of buprenorphine (0.03mg/Kg) was administered. Mice were weighed twice a week and ^1^H MRI was performed at 7T. Anaesthesia was induced using 3% isoflurane delivered in oxygen (0.5l/min) and maintained at 1-2%. Core body temperature was maintained using a thermo-regulated water-heated blanket or warm air blower. Following optimization of the magnetic field homogeneity, a rapid acquisition with relaxation enhancement (RARE) T2-weighted sequence (repetition time (TR) = 4500ms, effective echo time (TEeff) = 36ms, in-plane resolution 98μm × 98μm, 1mm thick contiguous slices) was used for localisation and assessment of tumours.

### Availability of models

Models may be requested, subject to available stocks, from the originating laboratories. For SU- cells, please contact Michelle Monje; for HSJD- cells, please contact Angel Montero Carcaboso; for OPBG- cells, please contact Maria Vinci. ICR-, BH-, DUB- and QCTB-cells are available from the CRUK Children’s Brain Tumour Centre of Excellence Cell Line Repository, which includes further information on growth conditions and morphology (crukchildrensbraintumourcentre.org)

### Data accessibility

All newly generated sequencing data have been deposited in the European Genome-phenome Archive (www.ebi.ac.uk/ega) with accession number EGAS00001004496 (sequencing) or ArrayExpress (www.ebi.ac.uk/arrayexpress/) with accession numbers E-MTAB-9297 (Illumina EPIC methylation arrays) and MTAB-9298 (Illumina 450k methylation arrays). Curated gene-level mutation, copy number, and expression data are provided as part of the pediatric-specific implementation of the cBioPortal genomic data visualisation portal (pedcbioportal.org).

## ACKNOWLEDGEMENTS

We acknowledge the patients and families whose lives have been impacted by these diseases and who have provided tissue for these studies. This work was supported by the INSTINCT network funded by The Brain Tumour Charity, Great Ormond Street Children’s Charity and Children with Cancer UK, the CRIS Cancer Foundation, Abbie’s Army, and Cancer Research UK. We further acknowledge funding from the Italian Ministry of Health, from Progetto Heal Onlus and Fondazione AIRC 5×100, 2018 Project code 21147. The authors acknowledge NHS funding to the National Institute for Health Research Biomedical Research Centre at The Royal Marsden and the ICR, and Experimental Cancer Medicines Centre (ECMC) and Royal Marsden Cancer Charity funding to the Paediatric Drug Development Team at The Royal Marsden and the ICR.

## AUTHOR CONTRIBUTIONS

DMC, ST, AM, MV and CJ conceived the study and wrote the manuscript. DMC, KK, LB, MC, NGO, KRT, MM, AMC and MV established *in vitro* models. ST, AM, EI, YG, PP, MH and CJ carried out molecular profiling and/or analysis. DMC, AM, HNP, RR, KK, LB, JFS, MC, AB, EFP, MV, CJL and CJ carried out drug screening, validation and/or analysis. DMC, KK, JKRB, VM, MF, KRT, SPR and MV carried out intracranial implantations and characterisation. CC, BZ, RB, AJM, BD, SS, SH, LVM, FC, HCM, SJV, SA-S, LRB, RJ, JC, MF, DC, JC, JP, FdB, AM, AC, MM, ASM, and TEGH provided primary patient material and annotation.

