## Supplementary Figures for "Drug screening linked to molecular profiling identifies novel dependencies in patient-derived primary cultures of paediatric high grade glioma and DIPG"

### SUPPLEMENTARY FIGURES AND TABLES

**Supplementary Table S1** – *List of patient-derived pHGG/DIPG cell cultures and basic phenotypic and molecular annotation.*

**Supplementary Table S2** – *Detailed molecular data for patient-derived derived pHGG/DIPG cell cultures.* (A) Short tandem repeat profiles. (B) Somatic mutations, including selected single nucleotide variants, indels, gene fusions, amplifications (AMP) and homozygous deletions (HOMDEL). (C) Methylation profiling – Heidelberg classifier scores.

**Supplementary Table S3** – *Drug screening data for patient-derived pHGG/DIPG cell cultures.* (A) Small screen SF50. (B) Small screen AUC. (C) Small screen Z score. (D) Large screen SF50. (E) Large screen AUC. (F) Large screen Z score.

**Supplementary Figure S1**

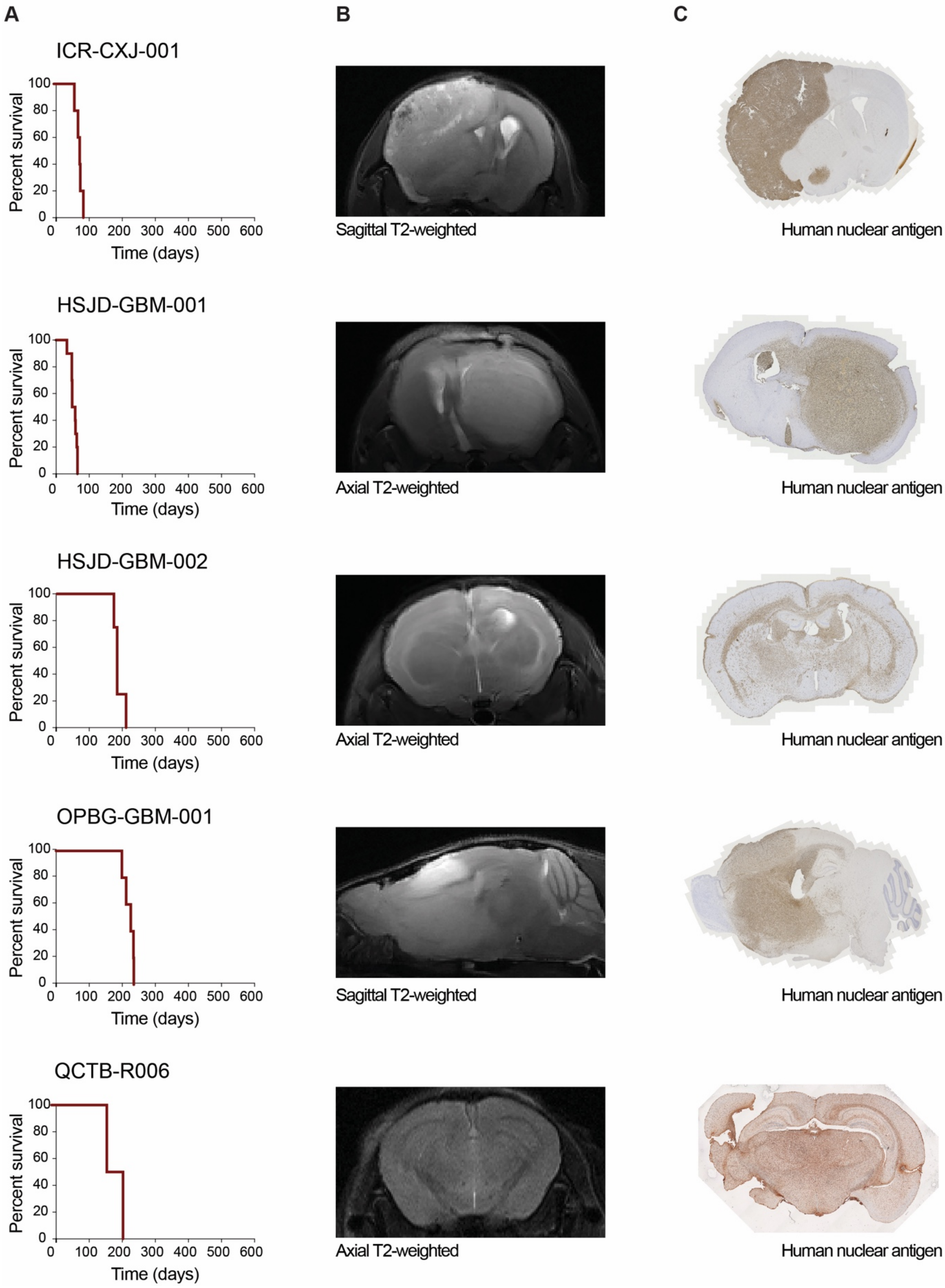

### Supplementary Figure S1

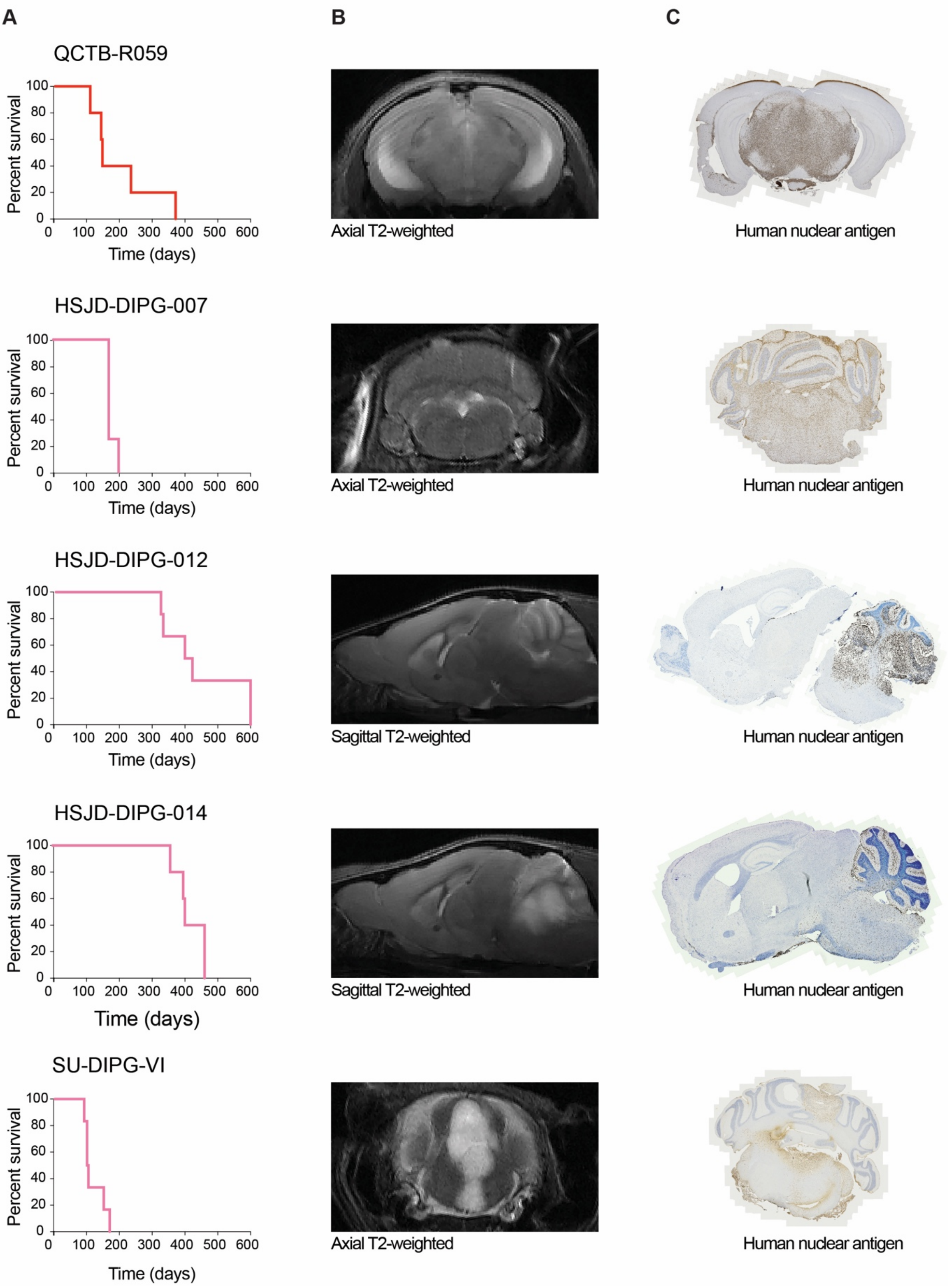

**Supplementary Figure S1** – Patient-derived orthotopic xenografts of pHGG/DIPG. (A) Survival curves for indicated models, with survival (y axis) plotted against time in days (x axis). (B) $T^2$-weighted MRI of indicated models, either sagittal or axial planes as labelled. (C) Human nuclear antigen staining of indicated models, either sagittal or axial sections as per MRI. Dark red: hemispheric HGG ICR-CXJ-001, HSJD-GBM-001, HSJD-GBM-002, OPBG-GBM-001, QCTB-R006; red: thalamic QCTB-R006; pink: pontine HSJD-DIPG-007, HSJD-DIPG-012, HSJD-DIPG-014, SU-DIPG-VI.

### Supplementary Figure S2

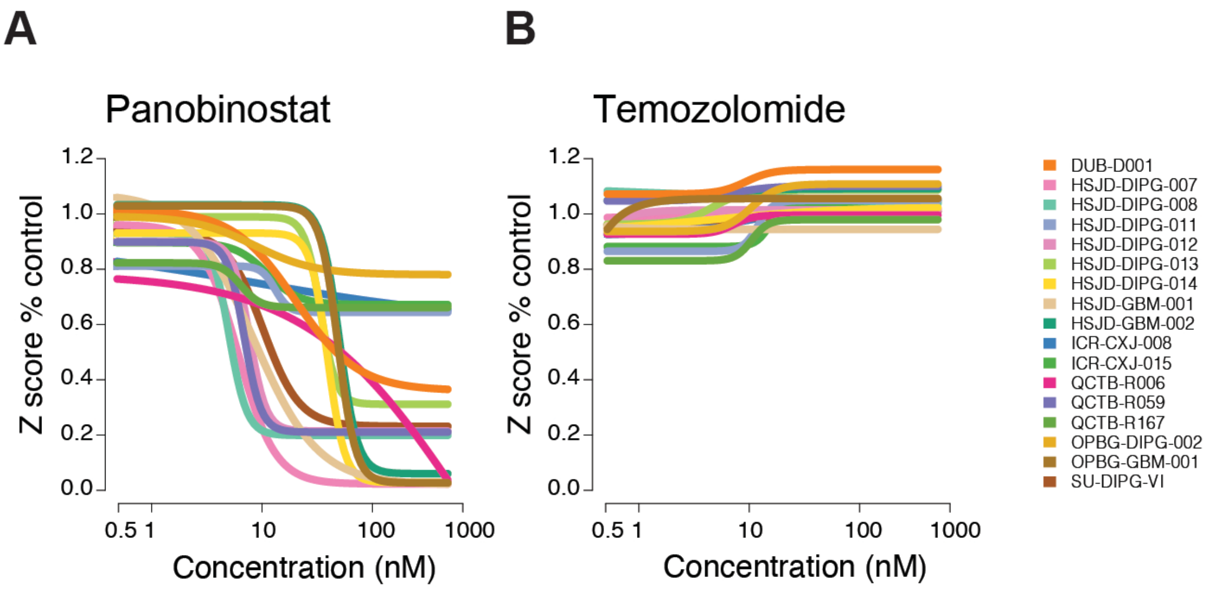


**Supplementary Figure S2** – *Drug screen data for commonly used agents in pHGG/DIPG.* (A) Screen data for the HDAC inhibitor panobinostat in the panel of pHGG/DIPG cultures, coloured according to the key provided. Concentration of compound is plotted on a log scale (x axis) against Z score plotted as a percentage of control (Z score POC) (y axis). (B) Screen data for the alkylating agent temozolomide in the panel of pHGG/DIPG cultures, coloured according to the key provided. Concentration of compound is plotted on a log scale (x axis) against Z score plotted as a percentage of control (Z score POC) (y axis).

### Supplementary Figure S3

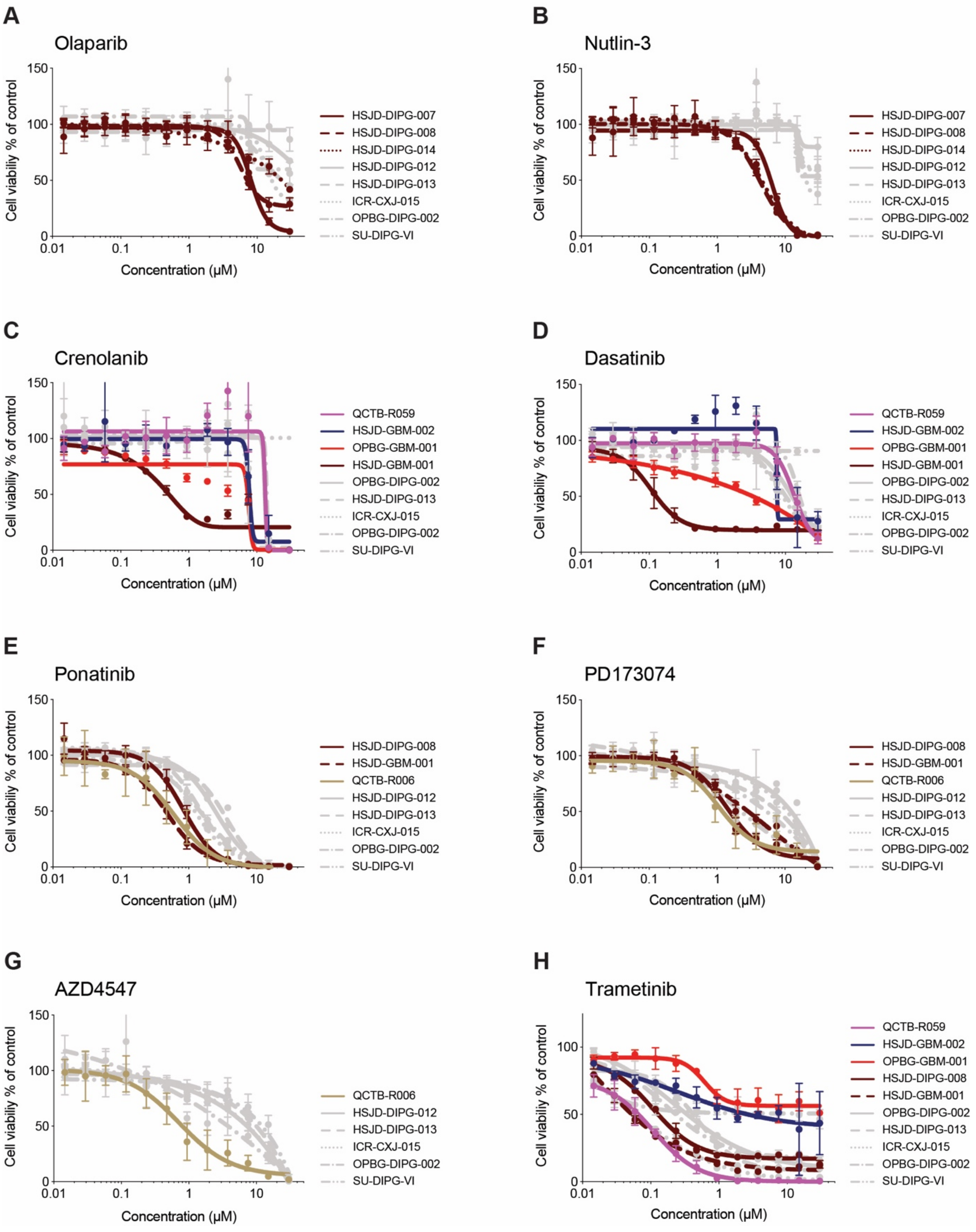


**Supplementary Figure S3** – *Validation dose-response curves for selected screening hits*. Validation dose-response curves for a variety of inhibitors in selected pHGG/DIPG cell cultures identified as sensitive in the initial screen (coloured, as below), and a mini-panel of less-sensitive controls (grey). (A) Olaparib. Dark red, *PPM1D*-mutant cells. (B) Nutlin-3. Dark red, *PPM1D*-mutant cells. (C) Crenolanib. *PDGFRA* mutations - dark red, sensitive; red, insensitive; dark blue, resistant; pink, wild-type amplification. (D) Dasatinib. *PDGFRA* mutations - dark red, sensitive; red, insensitive; dark blue, resistant; pink, wild-type amplification. (E) Ponatinib. Dark red, sensitising *PDGFRA* mutations; gold, *FGFR1*-overexpressing cells. (F) PD173074. Dark red, sensitising *PDGFRA* mutations; gold, *FGFR1*-overexpressing cells. (G) AZD4547. Gold, *FGFR1*-overexpressing cells. (H) Trametinib. *PDGFRA* mutations - dark red, sensitive; red, insensitive; dark blue, resistant; pink, wild-type amplification. Concentration of compound is plotted on a log scale (x axis) against cell viability (y axis). Mean plus standard error are plotted from at least n=3 experiments.
